# *fpocketR*: A platform for identification and analysis of ligand-binding pockets in RNA

**DOI:** 10.1101/2025.03.25.645323

**Authors:** Seth D. Veenbaas, Simon Felder, Kevin M. Weeks

## Abstract

Small molecules that bind specific sites in RNAs hold promise for altering RNA function, manipulating gene expression, and expanding the scope of druggable targets beyond proteins. Identifying binding sites in RNA that can engage ligands with good physicochemical properties remains a significant challenge. *fpocketR* is a software and framework for identifying, characterizing, and visualizing ligand-binding sites in RNA*. fpocketR* was optimized, through a comprehensive analysis of currently available RNA-ligand complexes, to identify pockets in RNAs able to bind small molecules possessing favorable properties, generally termed drug-like. Here, we demonstrate multiple, complex, uses of *fpocketR* to analyze RNA-ligand interactions and novel pockets in small and large RNAs, to assess ensembles of RNA structure models, to identify pockets in dynamic RNA systems, and to evaluate the shapes of RNA pockets. *fpocketR* performs best with RNA structures visualized at atomistic resolution but also provides useful information with lower resolution structures and computational models. *fpocketR* is a powerful, ligand-agnostic tool for discovery and analysis of targetable pockets in RNA molecules.

## INTRODUCTION

RNA molecules are essential regulators of gene expression, template protein synthesis, and carry out diverse additional cellular functions.^1,2^ As a result, RNAs lie upstream of nearly all biology. The vast scope of biological functions that can be modulated by altering RNA structure, translatability, or intermolecular interactions makes RNA an enticing target for small-molecule ligands.^3–8^ The field has made progress in targeting RNA with a few successful human-devised small molecules, currently limited to linezolid, an antibiotic that binds the ribosome,^9^ and risdiplam and branaplam, splicing modifiers that bind pre-messenger RNA.^10^ Better tools for assessing RNA ligands and their binding sites would facilitate exploiting the full potential of targeting RNAs with small molecules.

Much like proteins, RNA molecules routinely fold to form (base-paired) secondary structures and occasionally form higher-order tertiary structures. Small molecules can preferentially engage pockets formed within these complex tertiary RNA structures.^4,11,12^ However, physicochemical features of RNAs differ substantially from proteins: RNAs are generally more polar, more flexible, and more dynamic.^13–15^ These differences complicate the direct transfer of protein-based tools and heuristics to RNA. In addition, relatively few, roughly only 50,^12^ distinct RNA-ligand complexes have been visualized in which the ligand has good physicochemical properties (colloquially called drug-like). As a result of these two challenges, platforms for reliably identifying pockets in RNA that bind drug-like ligands are underdeveloped.

Foundational prior investigations have categorized small molecule interactions across diverse RNAs, based on comprehensive literature mining,^16,17^ on large-scale experimental binding measurements,^18,19^ on using available protein-focused software,^20,21^ and on analysis of high-resolution structure databases.^21,22^ These studies reinforce the idea that RNA motifs with complex structures are best able to form ligand-binding pockets, but have not focused on ligands with good physicochemical properties. Machine learning-based methods such as RLBind^23^ and MultiModRLBP^24^ were recently developed to predict the nucleotides in an RNA-ligand binding site. While powerful, these approaches typically do not return detailed geometric and physicochemical descriptors of complete binding pockets, limiting their ability to characterize ligand-binding environments in three dimensions. Finally, docking programs including AutoDock Vina,^25^ Dock 6,^26^ rDock,^27^ and RLDOCK^28^ have been widely applied to RNA.^29^ Docking requires a candidate ligand(s) and usually relies on generic pocket-finding routines to define the search space. Docking program strengths lie in pose prediction and affinity estimation, but do not address the complementary challenge of identifying which RNAs and which pockets are inherently capable of binding drug-like ligands in a ligand-agnostic manner. Overall, assessing RNA-ligand interactions is a rapidly growing area; nonetheless, characterization of RNA motifs specifically able to bind drug-like ligands remains incomplete.

We previously demonstrated that the open-source, geometry-based protein pocket finding software *fpocket*^30,31^ could be adapted to identify RNA-binding pockets through parameter optimization. Our optimization was guided by a curated training library of RNAs in complex with drug-like ligands. In a benchmark, we were able to detect all known ligand-binding sites in a dataset of 32 RNA-ligand complexes, selected for containing drug-like ligands, and we improved the positive predictive value from 19% for *fpocket* to 78%.^12^ We built on this optimization of *fpocket* and developed a pocket finding package specifically for RNA called *fpocketR*.^12^ Several major conclusions, unique to this work, emerged from the original development of *fpocketR*.^12^ First, *fpocketR* reliably detects pockets capable of binding ligands with good physicochemical properties. Second, *fpocketR* detected many novel, likely targetable, pockets in RNAs, an ability that was validated experimentally. Third, complex secondary structures, especially multi-helix junctions and pseudoknots, are an order of magnitude more likely to form pockets able to bind drug-like ligands than simpler motifs like RNA bulges and loops.

In this study, we now explore and validate multiple extended and more complex uses for *fpocketR* by characterizing pockets in large cryo-EM structures, computationally modeled RNA ensembles, and dynamic RNA systems. We further evaluate the geometric features consistent with forming RNA pockets for drug-like ligands. We anticipate that *fpocketR* will be a broadly useful ligand-agnostic framework for identifying and characterizing ligand-binding pockets in diverse RNA molecules.

## RESULTS AND DISCUSSION

### fpocketR workflow

*fpocketR* is a straightforward and streamlined analysis pipeline for identifying, visualizing, and characterizing pockets and ligands in RNA structures via a well-documented command-line interface. The pipeline is executed with a single input containing a Protein Data Bank (PDB) accession code or a locally stored RNA structure file (**Fig. 1A**). *fpocketR* also accepts RNA secondary structure drawing templates in multiple formats, including NSD,^32^ VARNA,^33^ and R2DT.^34^

**Figure 1.**
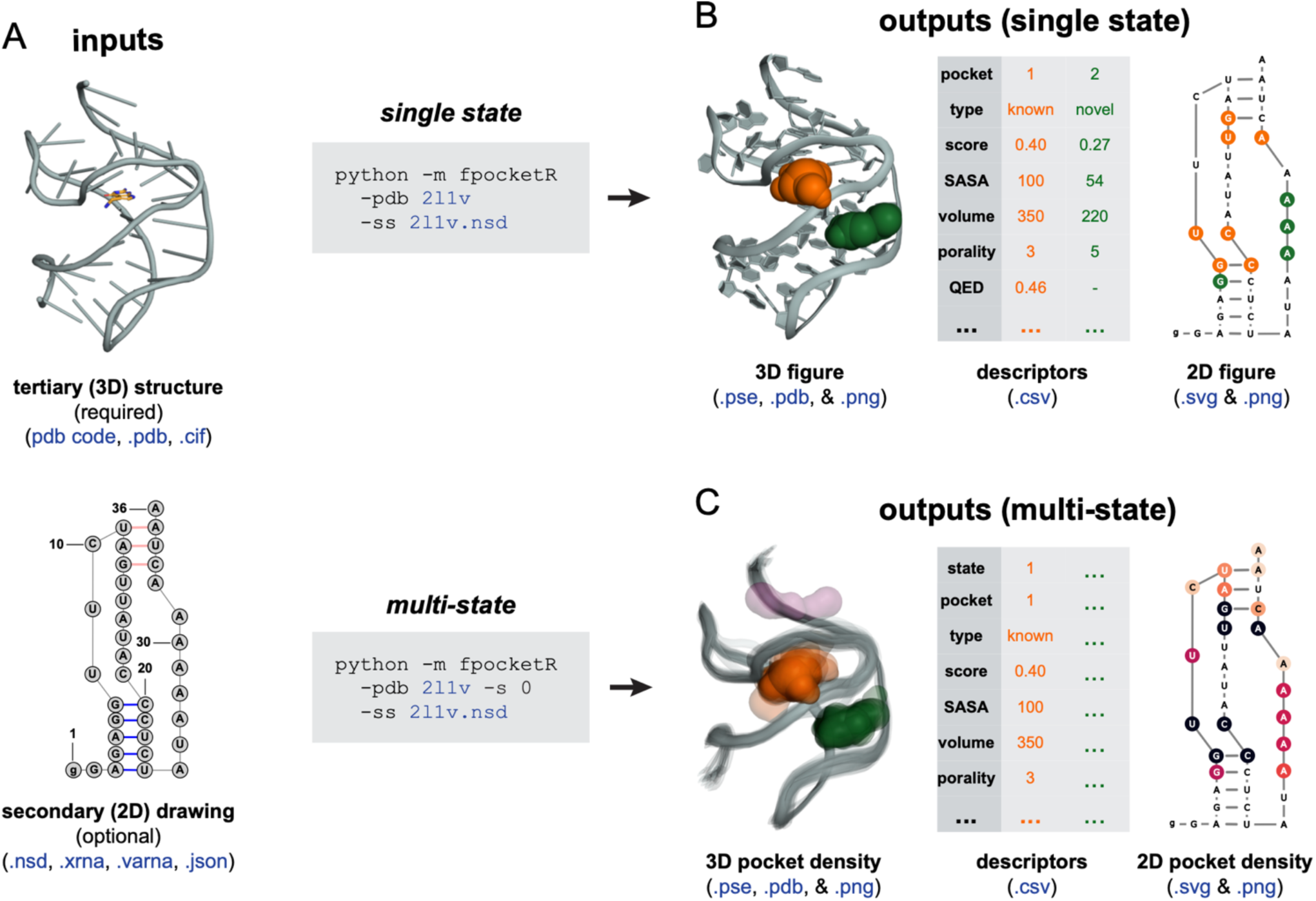
Pocket finding workflow for *fpocketR*, visualized with the class I preQ1 riboswitch (PDB: 2l1v). (A) Structural and command line inputs with accepted file formats. (B) Single-state analyses. Pockets are visualized as colored spheres and nucleotides that form pockets are highlighted in secondary structures. (C) Multi-state analyses. Map of tertiary structure pocket density displays all states of an RNA as a transparent backbone and pockets. In the map of pockets in secondary structure space, the intensity of colored nucleotides correlates with the frequency of participation in pockets.

*fpocketR* runs the *fpocket* algorithm with optimized parameters,^12^ characterizes pocket and ligand properties, ranks identified pockets by ligandability, and produces comprehensive output files. Outputs include a three-dimensional figure (and *pymol* session file) visualizing all pockets in the RNA tertiary structure, a detailed description of the properties of each pocket and ligand (**SI Table 2**), and a figure mapping the nucleotides that form each pocket onto an RNA secondary structure template (**Fig. 1B** and **SI Table 3**).

The behavior and features of *fpocketR* can be modified with optional arguments. A powerful feature of *fpocketR* enables analysis of multiple states in an RNA ensemble. Users can provide RNA ensembles from any source, including experimentally-determined structures, molecular dynamics simulations, or computational structure models. The multi-state analysis generates images that visualize the density of pocket formation in both tertiary and secondary structure formats (**Fig. 1C** and **SI Table 4**). Pocket finding can further be customized by manually adjusting pocket finding parameters and specifying which RNA chain(s) and ligand to analyze.

### Identification of pockets in large RNAs

Most available non-ribosomal RNA-ligand complexes involve short RNAs (< 200 nt), and many are riboswitches or simple aptamers.^22^ *fpocketR* was primarily trained on these small RNAs. However, application of cryo-electron microscopy (cryo-EM) has begun to expand the variety and sizes of solved RNA tertiary structures.^35–37^ We therefore examined the ability of *fpocketR* to detect and characterize pockets in large RNA structures. For the following examples, we emphasize that *fpocketR* has been optimized to selectively find pockets capable of binding drug-like ligands.^12^

We identified pockets in three large, RNA structures solved by cryo-EM. The first structure is a five-helix panel (544 nt), designed as an RNA origami scaffold (PDB 7ptq).^38^ The second structure is a large RNA (374 nt), engineered to include the small-molecule aptamers for Broccoli and Pepper, that bind the ligands DFHBI-1T and HBC620, respectively, configured as a Förster resonance energy transfer pair (7zj4).^39^ The third structure is a fungal group I intron bound to a pyrimidine small-molecule inhibitor (9mqt).^40^ Among the three RNAs, *fpocketR* identified all known pockets and seven novel pockets (**Fig. 2**). For the seven novel pockets, four were formed by pseudoknots (PK), one was formed at the interface (IN) of two helices, one by a multi-helix junction (MHJ), and one in the G-quadruplex (G4) of the Broccoli aptamer, adjacent to the DFHBI-1T binding site. The known pockets in the Broccoli-Pepper construct and the in the group I intron overlapped exactly with the ligand binding sites for DFHBI-1T, HBC620, and catalytic active site containing the pyrimidine inhibitor. This analysis, first, demonstrates that *fpocketR* detects high-quality binding pockets across a diverse range of RNA sizes and structure classes, despite the limited scope of its training dataset. Second, this analysis reinforces the idea that pockets able to bind drug-like ligands tend to form selectively in regions of complex local tertiary structure rather than in simple (helix and budged) RNA structures (**Fig. 2**).^12^

**Figure 2.**
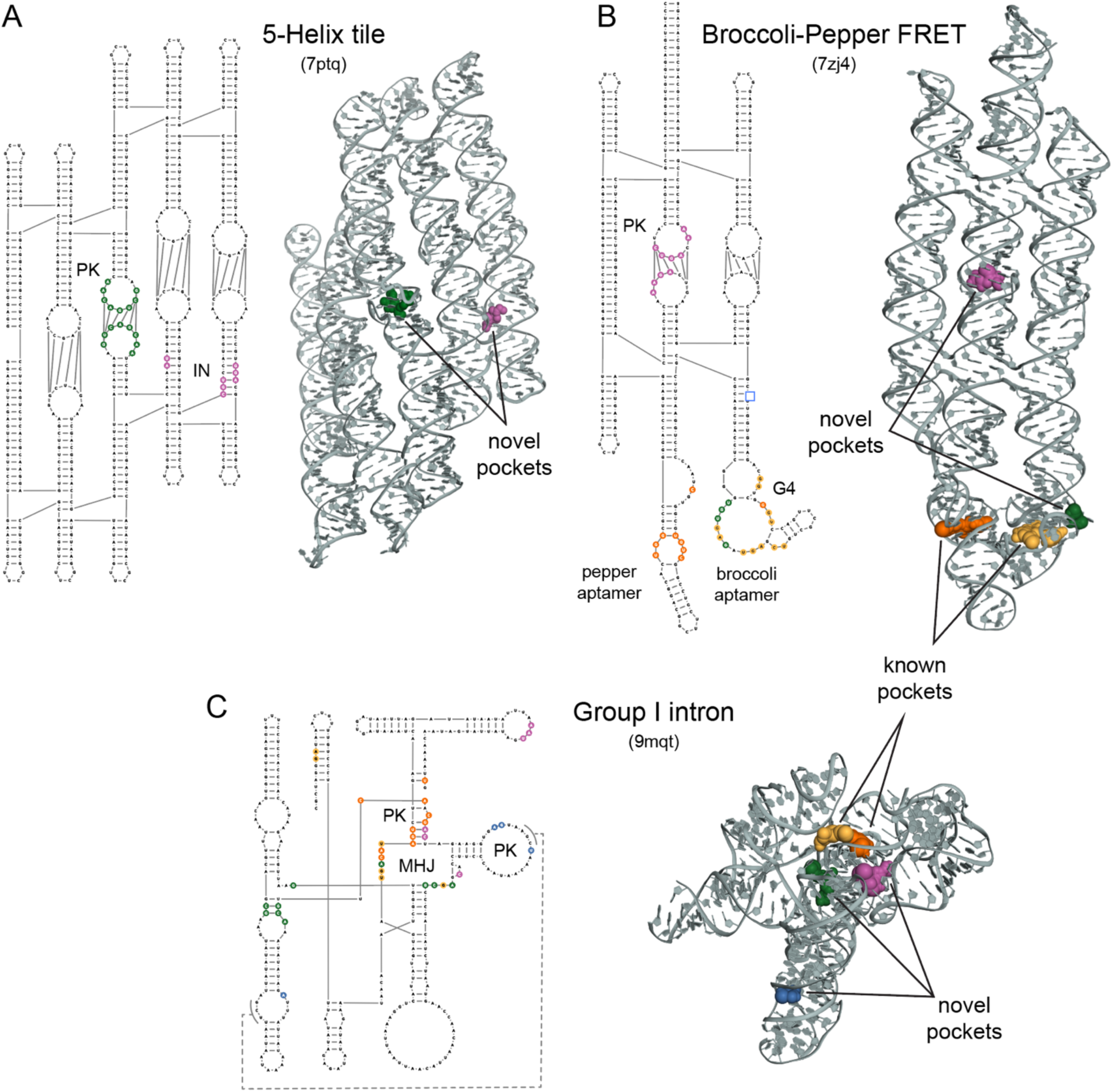
Pocket detection in large RNAs. Pockets detected in (A) a five-helix RNA origami scaffold (544 nts)^38^, (B) an RNA engineered to include Pepper (orange) and Broccoli (yellow) ligand-binding aptamers (374 nts)^39^, and (C) a fungal group I intron bound to a novel pyrimidine inhibitor (332 nts).^40^ RNA structures that form pockets are labeled as pseudoknot (PK), interface (IN), G-quadruplex (G4) or multi-helix junction (MHJ).

### Identification of pockets in low-resolution and modeled structures

Many RNA tertiary structures are solved at modest resolutions. Until recently,^37^ RNA-only structures solved using cryo-EM had resolutions in the 4 to 10 Å range.^36^ Modeling of RNA structure using physics-based or machine learning methods is rapidly improving, but still often exhibits large deviations from accepted structures.^41,42^ We investigated whether low-resolution and modeled structure ensembles can inform the ligandability of an RNA.

*fpocketR* detects one pocket in a high-resolution (2.3 Å) crystal structure of the class I type III preQ1 riboswitch (PDB 8fza),^43^ correctly identifying the ligand binding site for the preQ1 ligand (**Fig. 3A**). We applied *fpocketR* to 153 models of the same riboswitch submitted to the CASP15 evaluation.^41^ The models varied widely in accuracy, compared to the accepted structure, with root mean square deviations (RMSDs) between 2 and 28 Å and template modeling scores (TMscores) between 0.08 and 0.43. Of these models, 25% contained a pocket that overlapped the known preQ1 ligand binding site, 33% contained a pocket or pockets that did not overlap with the known ligand binding site, and 42% contained no pocket (**Fig. 3A**). We then compared the accuracy with which *fpocketR* detected the known ligand binding site in each model relative to the structural quality of the model. There is a modest correlation between pocket finding performance and the structural quality metrics RMSD and TMscore (**Fig. 3B**). The TMscore metric was the most informative, but still only modest, predictor of RNA ligandability, with one-half (48%) of the models with a TMscore over 0.26 containing a pocket overlapping the ligand binding site (**Fig. 3C**).

**Figure 3.**
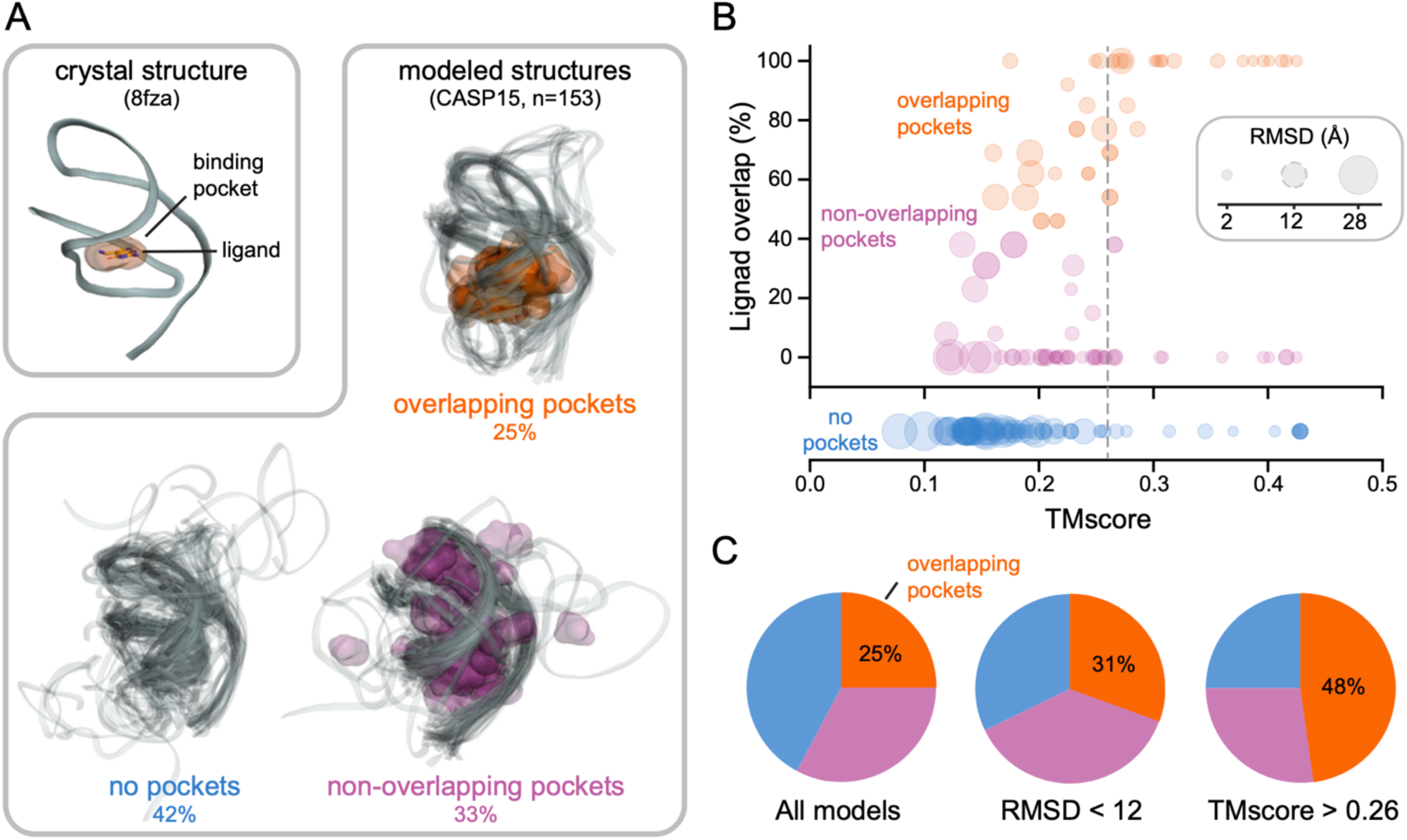
Pocket detection performance in modeled RNA structures. (A) Pockets detected in a reference structure (2.3 Å resolution; PDB 8fza)^43^ and in 153 models of the class I type III preQ1 riboswitch, from the CASP15 exercise.^41^ (B) Relationship between pocket performance (ligand overlap) and structural quality metrics, RMSD and TMscore. Dashed gray circular (RMSD) and vertical (TMscore) lines represent quality thresholds that produced the best selection for pockets overlapping the known ligand binding site. (C) Distribution of models with overlapping (orange), non-overlapping (pink), or no pocket (blue) for the indicated threshold of RMSD and TMscore.

We previously showed that *fpocketR* identified the SAM ligand binding site in a subset of states for a ligand-free ensemble of the SAM-IV riboswitch, determined by cryo-EM (PDB 6wql).^12,44^ We generated a 20-model ensemble of the SAM-IV riboswitch using trRosettaRNA, a successful machine-learning-based RNA modeling program,^41,45^ and then identified pockets in the structures of the resulting ensemble using the multi-state analysis mode from *fpocketR*. Pockets in the cryo-EM and modeled ensembles both clustered in the same two regions of the SAM-IV riboswitch: in the SAM ligand binding site and in the PK-1 pseudoknot (PK) (**Fig. 4**).

**Figure 4.**
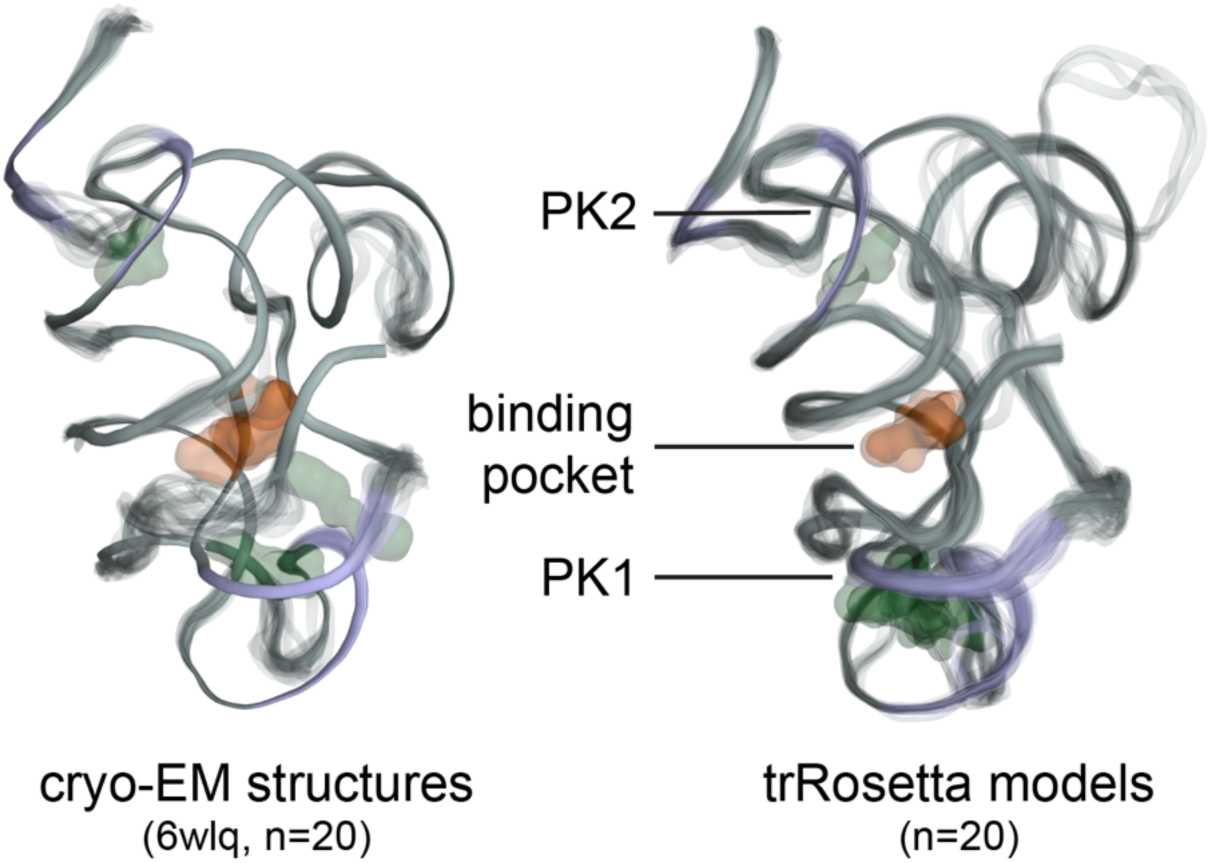
Pocket detection in multi-state ensembles. Map of pocket densities for ensembles of the SAM-IV riboswitch visualized by cryo-EM (PDB 6wlq)^35^ or generated by trRosettaRNA ^45^ modelling.

Overall, higher-accuracy RNA structures improve the accuracy of pocket finding with *fpocketR*. The performance of *fpocketR* with modeled structures for the preQ1 and SAM-IV riboswitches indicates that RNA modeling software can produce RNA structures with sufficient quality to assess the ligandability of the RNA.

### Identification of transient pockets in dynamic RNA structures

Changes in RNA structure and RNA dynamics are often critical for cellular function because structural alterations allow RNAs to respond to their environment and to interact with new partners.^46–48^ During translation, the ribosome undergoes large-scale conformational changes that include (*i*) rotation of the small and large ribosomal subunits in a rachet-like mechanism and (*ii*) swiveling of the head domain in the small ribosomal subunit.^49,50^ These large-scale movements enable concerted movement of messenger and transfer RNAs. Several antibiotics, including spectinomycin, neomycin, and Hygromycin B, inhibit translocation by stabilizing transient conformational states of the ribosome.^51^

We used *fpocketR* to search for pockets in ribosomal RNAs across six conformational states of the *E. coli* 70S ribosome during translocation including: PRE-C (7n1p), PRE-H1 (7n2u), PRE-H2 (7n30), INT1 (7n2v), INT2 (7n2c), and POST (7n31) (**Fig. 5A**).^50^ We then identified (transient) pockets present in only a subset of conformational states. We identified two pockets at the inter-subunit bridge, B2a, a conserved region located at the interface of helix 69 of the 23S rRNA, helix 44 of the 16S rRNA, and D-stems of the A- and P-site tRNAs.^52^ The transient pockets at the B2a bridge are not present in the early stages of translocation (**Fig. 5B**) and only form after back rotation of the small ribosomal subunit in the INT2 and POST states (**Fig. 5C**). These transient pockets partially overlap with binding sites for the antibiotics thermorubin^52^ and macrocyclic peptides viomycin^53^ and capreomycin^54^ (**Fig. 5D**). These antibiotics all have large masses (avg ∼650 Da), a large number of hydrogen bond acceptors (mean 13) and donors (mean 11), and low drug-likeness scores (avg quantitative estimate of drug-likeness, QED = 0.08).^55^ The (only) partial overlap of the pockets identified using *fpocketR* with binding sites visualized for these antibiotics likely reflects that large macrocyclic compounds do not require deep binding pockets, in contrast to the small molecules used to train *fpocketR*.

**Figure 5.**
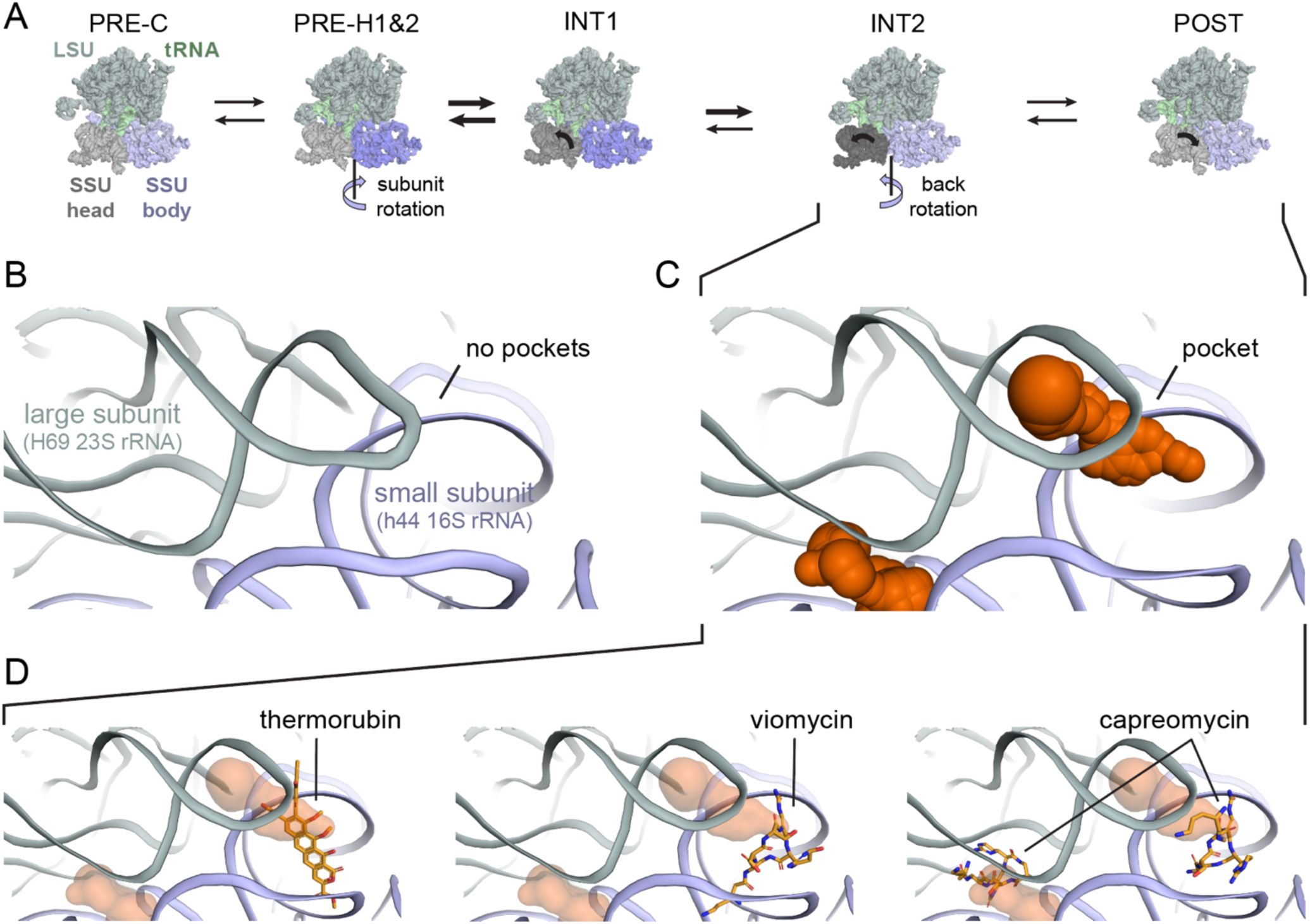
Identification of transient pockets at the interface of the large and small subunits of the bacterial ribosome, during translocation. (A) Overview of ribosome conformations at indicated stages of translocation.^50^ (B) Detailed view of the B2a inter-subunit bridge, which contains no pockets at early states of translocation. (C) Pockets in the B2a inter-subunit bridge formed between H69 and h44 in the INT2 and POST states of translocation. (D) Pockets detected at the B2a bridge region partially overlap binding sites for the antibiotics thermorubin (4v8a),^52^ viomycin (6lkq),^53^ and capreomycin (8ceu).^54^

Intriguingly, this analysis suggests that compact, drug-like ligands could be devised to bind these pockets, creating a path towards new antibiotics. This exercise demonstrates that *fpocketR* can identify regions that transiently form pockets in concert with RNA-mediated conformational changes. In principle, transient pockets can be targeted with small molecules, and understanding state-specific pocket formation could inform RNA-targeted drug mechanisms.

### Shapes of RNA pockets

Pockets in RNA, able to bind ligands with favorable physicochemical properties, are more polar and less hydrophobic compared to pockets in proteins.^12,20,21^ An important question is whether these differences influence the molecular shape of ligands that bind RNA. Molecular shape can be evaluated, independent of molecule size, using normalized ratios of principal moments of inertia (termed NPR values) to categorize molecular shapes broadly as rod-, disc-, or sphere-like.^56^ Diverse prior work has proposed that RNA ligands and pockets tend to be more rod-like and have fewer sphere-like shapes as compared to protein ligands and pockets.^17,20,21^

We calculated NPR values for the 376 pockets detected by *fpocketR* among non-redundant complexes in the Hariboss RNA-ligand database^22^ (<160 kDa; n = 364) and a single reference structure of the bacterial ribosome (PDB 7k00).^57^ This analysis used most available RNA-ligand complexes and includes ligands with a wide range of drug-likeness (QED 0.05 to 0.93). We emphasize that, by using *fpocketR*, we are focusing on pockets able to bind ligands with favorable physicochemical properties.^12^ RNA pockets appear to be mostly flat with shapes spanning the space from rod-like to disc-like. Both known (NPR1: 0.31, NPR1: 0.83) and novel (NPR1: 0.31, NPR2: 0.85) *fpocketR*-identified pockets have very similar distribution and average shape (**Fig. 6**). The distribution and average shape for both known and novel RNA pockets are strikingly similar to FDA-approved ligands (NPR1: 0.31, NPR2: 0.85).

**Figure 6.**
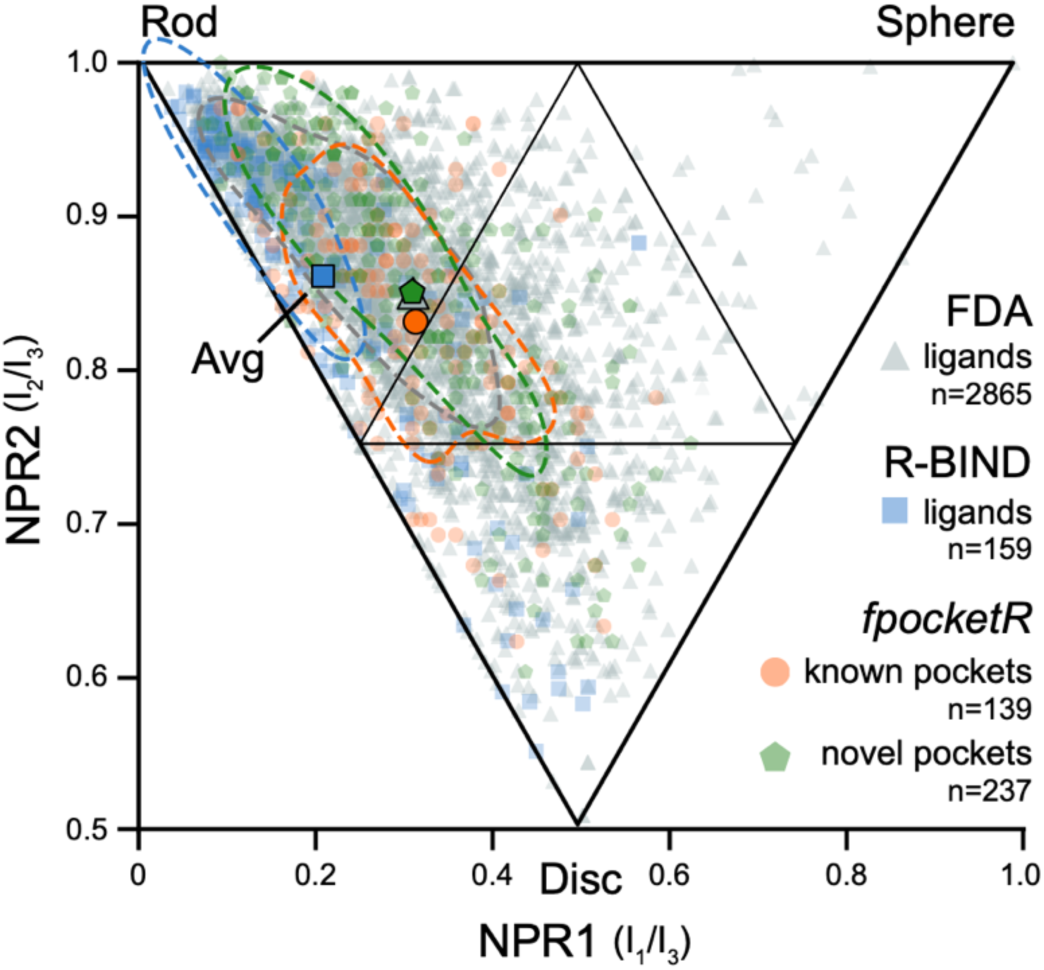
Comparison of shape-space for RNA-binding ligands versus RNA pockets. FDA-approved ligands (mostly protein binding), R-BIND ligands (exclusively RNA binding), and *fpocketR*-identified RNA pockets are shown. Normalized principal moment of inertia ratios (NPRs) for FDA-approved ligands (140-590 amu, n = 2865, grey triangles), R-BIND v2.1 small molecule ligands (n = 159, blue squares), and RNA pockets detected by *fpocketR* (known n = 139, orange circles; novel, n = 237, green pentagons). Each point represents the Boltzmann weighted average of NPRs for a single molecule using conformations within 3 kcal/mol of the lowest energy conformer. Large opaque symbols show the average shape for each category. Dashed lines represent the 50% contour line for each category. The distribution (Wasserstein) distances between FDA-approved ligands versus known RNA pockets, novel RNA pockets, and R-bind ligands are 0.038, 0.024, and 0.109, respectively; scale from 0-1, where 0 indicates identical distribution.

RNA pockets detected by *fpocketR* are not as rod-like as the RNA-associated ligands assessed in prior studies (**Fig. 6**). The difference in shape between pockets detected using *fpocketR* versus ligands examined previously^16,17^ likely reflects prior inclusion of long rod-like compounds that interact with RNA in the major and minor grooves and with sequence repeats. Ligands that bind in pockets are less rod-like than ligands that bind to repeat sequences.^58^ Together, these analyses suggest that RNAs contain pockets with a wide range of shapes able to bind ligands with equally diverse shapes (**Fig. 6**). The shape of RNA-targeted ligands is certainly important for selective engagement with a specific RNA pocket but, in bulk, the shapes of RNA-targeted and protein-targeted pockets and their ligands are broadly similar.

## CONCLUSIONS

*fpocketR* is a reliable ligand-agnostic platform that detects, characterizes, and visualizes high-quality ligand-binding pockets in RNA. In this study we demonstrate that *fpocketR* is broadly useful for complex systems involving RNA-ligand interactions and examine large RNAs, RNA ensembles, computational models, and dynamic conformational states. *fpocketR* identifies pockets in complex structural regions of large RNAs despite being trained on short (< 200 nt) RNA riboswitches and aptamers. The rules for the local structural environments that create ligand-binding pockets, capable of binding a drug-like ligand, are thus independent of macromolecular size. This study also provides further support that complex RNA structures – multi-helix junctions, pseudoknots, and other idiosyncratic tertiary structures – are the best RNA targets for drug-like small molecule ligands.

Unsurprisingly, pocket finding is most successful when structures are solved at high resolution. However, *fpocketR* also identified known pockets in computational models for the preQ1 and SAM-IV riboswitches examined here. These results demonstrate that RNA structures generated by physics-based and machine-learning modeling can be used to inform pocket detection and ligandability.

Many biologically important RNA targets are dynamic, and *fpocketR* provides dedicated and flexible tools for analyzing RNA ensembles. *fpocketR* identified transiently formed pockets at the interface between the small and large subunit rRNAs in the bacterial ribosome, which also overlapped with primary and secondary binding sites of translation-inhibiting antibiotics. The ability to identify RNA conformation states that selectively bind small-molecule ligands creates a powerful platform for guiding ligand (and drug) discovery and for defining binding and inhibition mechanisms.

*fpocketR* is optimized and validated to detect pockets capable of binding small molecules with favorable physicochemical properties, commonly termed drug-like.^12^ The shape-space of these RNA pockets closely matches the shape-space of FDA-approved small-molecule drugs, indicating that RNA-specific shape is probably not an important property to consider when curating libraries for RNA-targeted small molecules. Instead, chemical property differentiators other than shape should be the primary focus of RNA-focused ligand design.

*fpocketR* enables robust pocket detection for local RNA regions able to bind drug-like ligands, using both experimental and modeled RNA structures. The performance of *fpocketR* has been examined with and broadly validated for holo (with ligand), apo (without ligand), synthetic, dynamic, low-resolution, and computationally modeled RNA structures. We anticipate that *fpocketR* will provide diverse and thought-provoking information about the location and properties of ligand-binding pockets in RNA.

## METHODS

### Software availability

This study uses two newly developed software packages *fpocketR* and *MolMetrics*. *fpocketR*, used for pocket discovery and analysis, is freely accessed at https://github.com/Weeks-UNC/fpocketR. *fpocketR* requires a Unix-based operating system (for example, Linux or MacOSX). Windows users can run *fpocketR* via Windows Subsystem for Linux (WSL2) or a virtual machine. This repository includes usage documentation, example high-throughput workflows, and an explanation of all available features. *MolMetrics* generates multiple molecular conformers and calculates molecular descriptors and principal moments of inertia for small-molecule libraries via an efficient command line interface. *MolMetrics* is a cross-platform Python (3.7+) tool and is available at https://github.com/Weeks-UNC/molmetrics.

### Pocket finding

Pockets were identified, characterized, and visualized using *fpocketR* 1.2.0, which functions as a wrapper for *fpocket* v4.0.3.^30,31^ RNA tertiary and secondary structures were input using the *fpocketR* arguments --pdb and --nsd, respectively. Multi-state analyses were performed by setting the --state argument to 0 (all states).

### SAM-IV riboswitch 3D structure modeling

Structural models of the SAM-IV riboswitch were produced using the sequence from a reference structure^35^ (PDB: 6wlq) and secondary structure predicted by Fold from the RNAstructure package.^32^ These sequence and secondary structure files were then used to generate a multiple sequence alignment with rMSA^59^ using the NCBI nucleotide and RNAcentral databases.^60^ The multiple sequence alignment and predicted secondary structure were input to trRosettaRNA^45^ to generate 20 structural models. The resulting structures were aligned using PyMOL 3.0 (pymol.org, Schrödinger LLC) and saved as multiple states within a single (PDB) structure file.

### Multi-state analysis of the SAM-IV riboswitch

Ensembles for the SAM-IV riboswitch were analyzed using the *fpocketR* --state and --qualityfilter arguments. Pockets with a score less than 0.40 were omitted to facilitate direct comparison to previous studies.^12^

### R-BIND library

The RNA-targeted BIoactive ligaNd Database (R-BIND) (v2.1) molecules (downloaded on January 14, 2025) includes organic small-molecule probes reported in the literature through December 2021.^17^ The library was not filtered and contained 159 molecules.

### FDA-approved ligand library

FDA-approved (Phase 4) molecules^61^ (downloaded from CHEMBL on January 14, 2025) were filtered to exclude molecules with masses less than 140 amu or greater than 590 amu. The final library contained 2865 molecules.

### Hariboss RNA-ligand complex library

Hariboss RNA-ligand complexes^22^ (downloaded on January 14, 2025) were filtered to exclude redundant complexes and require RNAs be between 4 and 160 kDa (∼15 – ∼500 nts). A single high-resolution ribosome structure was added (PDB 7k00).^57^ The final library contained 365 RNA-ligand complexes (**SI Table 1**) which bind to both low and high QED score ligands (avg. QED = 0.35). Our curated library maximizes unique RNA-ligand complexes and contains approximately 160 RNA structures, many of which bind the same ligands (for example, TPP or SAM). *fpocketR* detected 139 known pockets, which selectively overlapped ligands with higher QED scores (avg. QED = 0.44), and identified 237 novel pockets. Notably, pockets detected in the final curated Hariboss library have a nearly identical average shape (NPR1: 0.30, NPR2: 0.84) to the pockets detected in the RNAs used to test and train *fpocketR* (NPR1: 0.31, NPR2: 0.84).^12^

### Principal moment of inertia analysis

Normalized principal ratios (NPRs) of principal moments of inertia were calculated for R-BIND and FDA-approved ligands using RDKit.^62^ Low-energy conformations (n=1000) were generated from SMILES strings using ETKDGv3. The lowest energy conformer was determined by optimizing geometries using the Universal Force Field. Boltzmann weighted average NPR values were calculated from all conformers within 3 kcal/mol of the lowest energy conformer. We note that the assessment of NPRs for a small molecule by RDKit does not account for the radius of the atoms in the molecule. In contrast, NPRs of RNA pockets calculated by *fpocketR* reflect a method that takes the radius of alpha cores into account.

*fpocketR* calculates the principal moments of inertia for pockets after converting each pocket from a cluster of alpha cores (alpha core = alpha sphere – 1.65 Å) into a single solid object with uniform density. The shape of RNA pockets detected with *fpocketR* is shifted to-0.07 on the NPR1 axis, relative to values provided by RDKit. We therefore normalized NPR1 values to the left diagonal (running between rod-like and disc-like vertices) to normalize NPR values calculated for pockets and ligands.

Contour lines were generated using kernel density estimation at a 50% density level, and joint Wasserstein distances were computed between bivariate distributions using the Python Optimal Transport Theory library,^63^ quantifying the multidimensional dissimilarity relative to the distribution of FDA-approved ligands.

## ACKNOWLEGEMENT

We are indebted to Patrick Irving for providing thoughtful feedback on this work. This study was supported by grants from the US National Science Foundation (MCB-2027701 to K.M.W.) and National Institutes of Health (R21 AG084970 to S.F. and K.M.W.).

## SUPPORTING INFORMATION

Supporting Tables (4), description of Supporting Dataset (1), and Archive of Pymol Session Files (1).

---

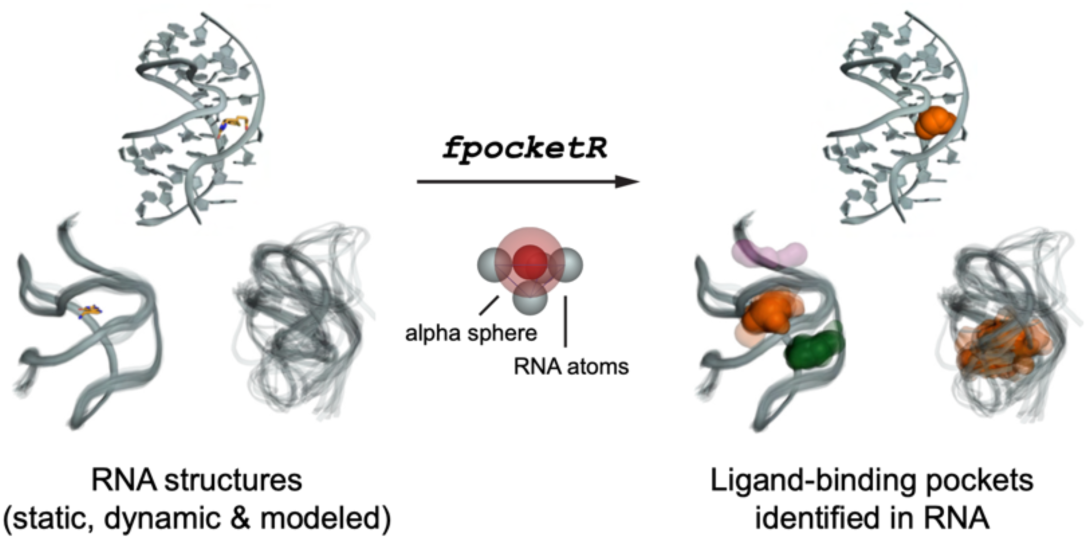

