## Supporting Information for "*fpocketR*: A platform for identification and analysis of ligand-binding pockets in RNA"

Supporting Tables (4), description of Supporting Dataset (1), and Archive of Pymol Session Files (1).

**SI Table 1.** PDB accession codes for non-redundant Hariboss RNA-ligand complex library (≤ 160 KDa).

| | | | | | | | | | |
|---|---|---|---|---|---|---|---|---|---|
| 1aju | 2au4 | 2tra | 3ski | 4q9r | 5v3f | 6e84 | 6up0 | 7k00 | 8d2b |
| 1am0 | 2be0 | 2yie | 3skz | 4qk9 | 5v9z | 6e8t | 6v9b | 7kuk | 8d5l |
| 1arj | 2bee | 3b4a | 3slm | 4qln | 5vcf | 6e8u | 6v9d | 7kum | 8d5o |
| 1eht | 2esi | 3b4b | 3slq | 4qvi | 5vci | 6fz0 | 6va2 | 7kun | 8eyu |
| 1ei2 | 2esj | 3bnq | 3suh | 4ts2 | 5vj9 | 6gzr | 6va3 | 7kuo | 8eyv |
| 1evv | 2et3 | 3d2g | 3sux | 4w90 | 5vjb | 6hag | 6va4 | 7kup | 8eyw |
| 1f1t | 2et4 | 3d2v | 3td1 | 4xwf | 5xi1 | 6hbt | 6vui | 7kvu | 8f4o |
| 1f27 | 2et8 | 3d2x | 3tzr | 4y1j | 5z1h | 6hc5 | 6wzr | 7kvv | 8fza |
| 1fmn | 2f4t | 3dig | 3v7e | 4y1m | 5z1i | 6hmo | 6wzs | 7lne | 8gxb |
| 1fuf | 2f4u | 3dil | 3wru | 4yaz | 5z71 | 6jbf | 6xb7 | 7lnf | 8gxc |
| 1fyp | 2g5k | 3diq | 4e8k | 4yb0 | 5zei | 6jbg | 6xkn | 7lng | 8hb3 |
| 1i2y | 2gdi | 3dir | 4e8q | 4yb1 | 5zej | 6jji | 6xrq | 7mkt | 8hb8 |
| 1i9v | 2ho7 | 3ds7 | 4e8v | 4zc7 | 6az4 | 6lau | 6yl5 | 7oa3 | 8hba |
| 1j7t | 2hom | 3e5c | 4f8u | 4znp | 6bfb | 6lax | 6ylb | 7oaw | 8i3z |
| 1j8g | 2hoo | 3e5e | 4far | 5bjo | 6c64 | 6laz | 6ymi | 7oax | 8i43 |
| 1koc | 2hop | 3e5f | 4fau | 5bjp | 6c65 | 6n5o | 6ymj | 7rwr | 8i45 |
| 1kod | 2juk | 3egz | 4faw | 5btp | 6c8e | 6p2h | 6ymk | 7szu | 8i46 |
| 1lc4 | 2kd4 | 3f2q | 4gpx | 5c45 | 6c8k | 6pq7 | 6yml | 7td7 | 8i7n |
| 1lvj | 2kgp | 3f4g | 4gpy | 5dhb | 6c8l | 6q57 | 6ymm | 7tdc | 8r62 |
| 1mwl | 2ktz | 3f4h | 4jf2 | 5fjc | 6c8m | 6qiv | 7a3y | 7tzr | 8r63 |
| 1n7a | 2ku0 | 3fo4 | 4k31 | 5fk1 | 6c8o | 6t3n | 7d12 | 7tzt | 8r8p |
| 1n7b | 2kx8 | 3fo6 | 4k32 | 5fk4 | 6cb3 | 6tb7 | 7d7v | 7tzu | 8swg |
| 1nta | 2kxm | 3gca | 4kqy | 5fk6 | 6cc1 | 6tf0 | 7d7w | 7u0y | 8swo |
| 1ntb | 2l1v | 3ger | 4kzd | 5hbw | 6cc3 | 6tf1 | 7d81 | 7u87 | 8sx5 |
| 1o15 | 2l94 | 3gx2 | 4l81 | 5kpy | 6ck4 | 6tf2 | 7dwh | 7u88 | 8sx6 |
| 1q8n | 2lwk | 3gx3 | 4lvv | 5krg | 6ck5 | 6tf3 | 7e9e | 7u89 | 8sxl |
| 1raw | 2m4q | 3iwn | 4lvx | 5kx9 | 6db8 | 6tfe | 7edl | 7u8a | 8sy1 |
| 1tn2 | 2miy | 3k0j | 4lvy | 5lwj | 6dlr | 6tfg | 7edt | 7u8b | 8u5k |
| 1uts | 2mxs | 3k1v | 4lx5 | 5ob3 | 6dmc | 6u6j | 7elp | 7wib | 8u5p |
| 1uud | 2n0j | 3mij | 4lx6 | 5t83 | 6dn1 | 6u89 | 7eog | 7wif | 8u5t |
| 1uui | 2o3v | 3muv | 4nfo | 5ued | 6dn2 | 6u8f | 7eok | 7wii | 8u5z |
| 1xpf | 2o3w | 3mxh | 4nya | 5uee | 6dn3 | 6u8u | 7eol | 7ych | 8vaw |
| 1ykv | 2o3x | 3npq | 4nyb | 5ueg | 6e1s | 6uc7 | 7eom | 7yci | |
| 1yls | 2o3y | 3q3z | 4p20 | 5ux3 | 6e1u | 6uc8 | 7eon | 7zj4 | |
| 1yrj | 2oe5 | 3rkf | 4p5j | 5v0h | 6e1v | 6uc9 | 7eoo | 8cf2 | |
| 1zz5 | 2pwt | 3s4p | 4pdq | 5v0o | 6e1w | 6uej | 7eop | 8d28 | |
| 2a04 | 2qwy | 3sd3 | 4q9q | 5v1l | 6e81 | 6uet | 7fhi | 8d2a | |

**SI Table 2.** Description of all pocket characteristics generated by *fpocketR*.

| Characteristic | Description |
|---|---|
| Parameters | fpocket parameters used to find pockets. |
| Name | Structure name generated from input file. |
| PDB | First four characters of the inputted tertiary structure (typically the PDB accession code). |
| State | Conformational state from the tertiary structure file. |
| Type | **Known**: pocket overlaps with ligand. **Novel**: pocket does not significantly overlap with ligand. |
| Filter | **Pass**: pocket score above the quality filter threshold. **Fail**: pocket is not visualized in figures. |
| Pocket | Pocket ID number. |
| Score | † Pocket score from fpocket. |
| Drug_score | † Pocket drug score from fpocket. |
| a-sphere | † Number of alpha spheres in the pocket. |
| SASA | † Solvent accessible surface area. |
| Volume | † Volume of pocket (Å$^3$). |
| Hydrophobic _density | † Sum of all apolar alpha sphere neighbors divided by the # of apolar alpha spheres in the pocket. |
| Apolar_ a-sphere_ proportion | † Percentage, reflects the proportion of apolar alpha spheres among all alpha spheres in the pocket. |
| Hydrophobicity_score | † Residue-based hydrophobicity scale reported in Monera & al. (1995) *Prot Sci*, **1**, 319-329. |
| Polarity_score | † Measure the hydrophilicity character of the pocket. |
| PocketNT | List of all RNA residues that contact alpha spheres from the pocket. |
| Pocket_NPR1 | First normalized principal momental of inertia ratio ($I_1/I_3$) for the pocket. |
| Pocket_NPR2 | Second normalized principal momental of inertia ratio ($I_2/I_3$) for the pocket. |
| Pocket_shape | Description of pocket shape based on NPR: **rod-like**, **disc-like**, **sphere-like**, or **balanced**. |
| Ligand_ID | Chemical ID of the ligand assigned from the PDB Chemical Component reference Dictionary. |
| Pocket_overlap | Proportion of alpha sphere in the pocket that are within 3 Å of an atom in the ligand. |
| Ligand_overlap | Proportion of atoms in the ligand that are within 3 Å of an alpha spheres from the pocket. |
| Center_criteria | Minimum Euclidean distance between the geometric centers of the pocket and ligand. |
| QED_score | Ligand drug-likeness score reported in Bickerton,G.R. (2012) *Nat Chem*, 4, 90-98. |
| Ligand_NPR1 | NPR1 ($I_1/I_3$) for the ligand calculated using the conformation from the PDB (not optimized!). |
| Ligand_NPR2 | NPR2 ($I_2/I_3$) for the ligand calculated using the conformation from the PDB (not optimized!). |
| Ligand_shape | Description of ligand shape based on NPR: **rod-like**, **disc-like**, **sphere-like**, or **balanced**. |

† Characteristics calculated and documented by *fpocket* v4.0.3.

**SI Table 3.** Output files used and generated by *fpocketR*.

| Filename | Description |
|---|---|
| <PDB>clean_out/<ligand_id>_model.sdf | Analyzed ligand structure from the PDB. |
| <PDB>clean_out/pockets/pocket<#>_atm.pdb | RNA atoms that contact alpha spheres used to determine pocketNT. |
| <PDB>clean_out/pockets/pocket<#>_surf.obj | Pocket surface used to calculate NPR values. |
| <PDB>clean_out/pockets/pocket<#>_vert.pqr | Alpha spheres in pocket used to generate *_real_sphere.pdb. |
| <PDB>clean_out/<PDB>_out_pocket_characteristics.csv | Geometric and physicochemical properties of pockets. |
| <PDB>clean_out/<PDB>_2D.png | Figure of pockets in RNA secondary structure. |
| <PDB>clean_out/<PDB>_2D.svg | Editable figure of pockets in RNA secondary structure. |
| <PDB>clean_out/<PDB>_out_real_sphere.pdb | RNA structure with pockets. Alpha sphere radius in B factor column. |
| <PDB>clean_out/<PDB>_out_real_sphere.pse | RNA structure with properly scaled alpha spheres in pockets. |
| <PDB>clean_out/<PDB>_3D_<dpi>.png | Figure of pockets in RNA tertiary structure. |

**SI Table 4.** Output files used and generated by *fpocketR* multistate analysis.

| Filename | Description |
|---|---|
| state_tracker.txt | Stores last state analyzed used to resume interrupted analysis. |
| <PDB>_all_states_out_pocket_characteristics.csv | Geometric and physicochemical properties of pockets in all states. |
| <PDB>_2D_pocket_density.png | Figure of pockets density from all states in RNA secondary structure. |
| <PDB>_2D_pocket_density.svg | Editable figure of pockets density from all states in RNA secondary structure. |
| <PDB>_all_states_out_real_sphere.pse | RNA structure of all states with pockets depicted as transparent surfaces. |
| <PDB>_all_states_3D_<dpi>.png | Figure of pockets in all states of RNA tertiary structure. |

**SI Dataset 1.** Principle moment of inertia analysis.

Excel file containing identity and descriptors (including SMILES, QED scores, selected molecular descriptors, and NPR values) for the ligands and pockets used in the principal moment of inertia analysis (**Fig. 6**).

**Archive of Pymol Session Files.** Pymol session files showing images in Figures 2–5.
